# Long-Timescale Molecular Dynamics Reveal a Coordination-Biased Conformational Selection Mechanism for Sorcin Activation

**DOI:** 10.64898/2026.07.08.737357

**Authors:** Qiushi Ye, Ibrahim D. Boyenle, Holly Hemesath, Kathleen Joyce Carillo, Mithras Manssuri, Lei Zhang, Yanxin Liu

## Abstract

Sorcin is a dimeric penta-EF-hand Ca^2+^-binding protein that regulates intracellular Ca^2+^ homeostasis through Ca^2+^-dependent conformational activation and target recognition, and it has also been implicated in multidrug resistance in cancer. Although crystal structures have defined the apo inactive and Ca^2+^-bound active states of Sorcin, the transition pathways connecting these states and the conformational ensembles populated under each condition remain poorly understood. Here, we used long-timescale all-atom molecular dynamics simulations on Anton 3, totaling ∼90 μs, to define the Ca^2+^-coupled conformational landscape of dimeric human Sorcin at atomic resolution. Starting from the Ca^2+^-bound structure, we directly observed the transition from the active to the inactive state following Ca^2+^ removal, demonstrating that loss of Ca^2+^ coordination is sufficient to drive inactivation on the microsecond timescale. Simulations initiated from the Ca^2+^-bound crystal structure with retained ions unexpectedly revealed ultrafast Ca^2+^ dissociation and rebinding at all EF-hand sites, indicating weak intrinsic Ca^2+^ affinity and highly dynamic ion exchange. In complementary simulations initiated from the apo structure, Sorcin spontaneously sampled active-like conformations even in the absence of stable Ca^2+^ binding, supporting a conformational selection mechanism in which Ca^2+^ shifts the population toward pre-existing active states rather than inducing the transition de novo. Across all conditions, we also observed pronounced and persistent structural asymmetry between the two protomers, revealing that the Sorcin homodimer is dynamically heterogeneous despite its symmetric crystal structures. Together, these results support a coordination-biased conformational selection model for Sorcin activation, in which weak and rapidly exchanging Ca^2+^ binding stabilizes, rather than induces, the active state. This work provides a dynamic framework for understanding Sorcin function as a fast Ca^2+^ sensor and offers broader mechanistic insight into activation principles of EF-hand Ca^2+^-binding proteins.

## Introduction

Sorcin (<u>so</u>luble resistance-related <u>c</u>alcium-binding prote<u>in</u>) has emerged as a clinically relevant regulator in multiple disease contexts, particularly in cancer and neurodegenerative disorders(*1–6*). In cancer, Sorcin is frequently overexpressed and has been strongly associated with multidrug resistance(*7–9*). It modulates intracellular calcium signaling to promote cell survival and reduce apoptotic sensitivity(*4,5*). Elevated Sorcin levels have been observed in leukemia, breast cancer, and other malignancies(*9,10*). It contributes to chemotherapeutic failure and poor prognosis(*2*). In parallel, dysregulation of Sorcin has also been implicated in neurodegenerative diseases, including Alzheimer’s and Parkinson’s disease(*6*), where altered calcium homeostasis and protein misfolding are central pathological features. Sorcin’s ability to interact with key calcium-handling proteins and influence endoplasmic reticulum function positions it as a potential modulator of cell vulnerability and stress responses, linking its molecular behavior to broader disease mechanisms(*2,5*).

Functionally, Sorcin plays a central role in maintaining cellular calcium homeostasis through its activity as a penta-EF-hand calcium-binding protein(*1,11*). Upon binding Ca^2+^, Sorcin undergoes conformational changes that enable it to interact with and regulate a variety of target proteins, including the ryanodine receptor, L-type calcium channels, and SERCA pumps(*12–15*). Through these interactions, Sorcin helps fine-tune calcium flux between the cytosol and intracellular Ca^2+^ storing organelles, particularly within the endoplasmic and sarcoplasmic reticulum. This regulation is essential for processes such as excitation-contraction coupling(*14*), signal transduction(*16,17*), and cellular stress responses(*18*). The calcium-dependent activation and inactivation of Sorcin, therefore, represents a key molecular switch that links ion binding to functional output, underscoring the importance of understanding its activation mechanism at a structural and dynamical level.

Sorcin belongs to the penta-EF-hand family of Ca^2+^-binding proteins and exists as a homodimer composed of a Ca^2+^-binding C-terminal domain (CTD; also referred to as the calcium-binding domain) and a flexible glycine-rich N-terminal domain (NTD), as shown in Figure 1A(*19–21*). Crystal structures of the apo states (PDB ID: 4UPG) and the Ca^2+^-bound (PDB ID: 4USL) reveal substantial rearrangements within the CTD, particularly involving helices D and G(*20*). This Ca^2+^-induced conformational transition can be quantified by the angle between helices D and G, which increases from 57° in the apo inactive state to 78° in the Ca^2+^-bound active state (Figure 1B and 1C)(*20*). We recently completed the NMR backbone resonance assignments of full-length Sorcin in the apo state(*22*). However, Ca^2+^-induced Sorcin aggregation has prevented further investigation of its conformational dynamics in the Ca^2+^-bound state(*23*). Moreover, existing structural approaches remain limited in their ability to define transition pathways, quantify microscopic state populations, or resolve the extent of chain asymmetry within the Sorcin dimer. As a result, a comprehensive understanding of the dynamic conformational landscape underlying Sorcin activation remains incomplete.

**Figure 1.**
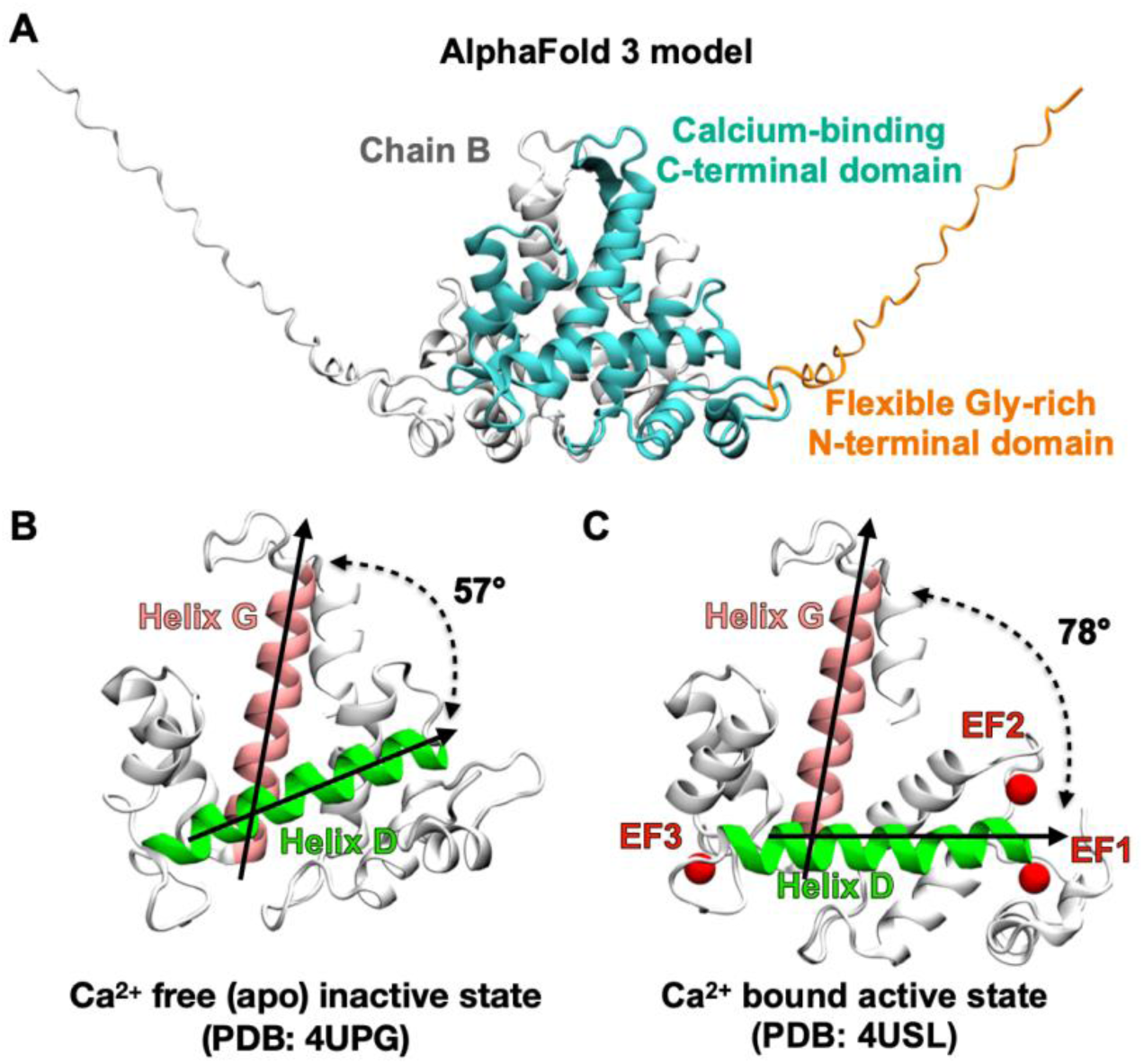
Atomic structure of human sorcin. (A) AlphaFold 3-predicted model of full-length human Sorcin in its dimeric form. Chain A is colored with the Gly-rich N-terminal domain (NTD) in orange and the calcium-binding C-terminal domain (CTD) in cyan. Chain B is colored in white. (B) Crystal structure of the Ca^2+^-free (apo), inactive state of Sorcin (PDB ID: 4UPG). The angle between helix D (green) and helix G (pink) is 57° in the inactive state. (C) Crystal structure of the Ca^2+^-bound, active state of Sorcin (PDB ID: 4USL). Bound Ca^2+^ ions in EF hands 1-3 are shown as red spheres. The angle between helix D (green) and helix G (pink) is 78° in the active state. Most of the flexible NTD were not resolved in the crystal structures.

Significant progress has been made in understanding Ca^2+^-induced Sorcin aggregation and its sensitivity to Ca^2+^(*23*) Our recent stopped-flow light-scattering measurements demonstrate that Ca^2+^-induced Sorcin self-association and aggregation are both reversible and cooperative(*23*). These light-scattering aggregation assays further reveal that Sorcin exhibits relatively low sensitivity to Ca^2+^, with activation occurring at concentrations close to 100 micromolar range(*23*). This apparent Ca^2+^ sensitivity is further modulated by Mg^2+^ and ionic strength(*23*). Together, these findings support an activation mechanism in which Ca^2+^ binding shifts Sorcin toward conformations with increased exposure of hydrophobic and target-recognition surfaces(*20,23*). In the absence of binding partners, Ca^2+^-activated Sorcin can also undergo transient self-association and aggregation(*23*). A detailed structural framework is therefore needed to connect these kinetic observations in aggregation assay to underlying molecular motions. Specifically, it remains unclear which conformational ensembles are populated under apo and Ca^2+^-bound conditions, how structural transitions proceed following Ca^2+^ binding or release, and how stable coordination is at individual EF-hand sites. Resolving these questions will be essential for linking macroscopic aggregation behavior to the microscopic dynamics that govern Sorcin activation.

In this study, we employed long-timescale molecular dynamics (MD) simulations performed on newly available Anton 3(*24*) to elucidate the mechanisms underlying Ca^2+^-dependent conformational transitions in Sorcin and to characterize the structural ensembles of its active and inactive states. Our simulations directly capture the transition from the Ca^2+^-bound active state to the apo inactive state upon Ca^2+^ release and reveal unexpectedly rapid Ca^2+^ binding and unbinding kinetics at EF-hand sites. By systematically modulating Ca^2+^ coordination through restraint schemes, we demonstrate that the active conformation is stabilized by Ca^2+^ binding, whereas loss of coordination drives the transition to the inactive state. Notably, we find that the active state can be spontaneously sampled even in the absence of Ca^2+^, indicating that Ca^2+^ binding shifts the conformational equilibrium rather than inducing the transition de novo. Furthermore, our simulations uncover pronounced structural asymmetry within the Sorcin homodimer, suggesting an additional layer of functional regulation. Together, these results support a conformational selection mechanism for Sorcin activation and provide a dynamic, atomistic framework for understanding its role as a Ca^2+^ sensor.

## Methods

### Atomic modeling of full length Sorcin

Atomic models of dimeric human Sorcin were constructed based on crystal structures representing the apo (PDB ID: 4UPG) and Ca^2+^-bound (PDB ID: 4USL) states(*20*). Structured regions were taken directly from the corresponding crystal structures (residues 30-198 for the apo state and residues 29-198 for the Ca^2+^-bound state). The remaining N-terminal residues (residues 2-29 for the apo state and residues 2-28 for the Ca^2+^-bound state) were modeled using AlphaFold3(*25*). The missing hydrogen atoms were generated using the psfgen plugin in VMD(*26*) with the CHARMM36m(*27,28*) topology file.

To investigate Sorcin inactivation and Ca^2+^ release-induced conformational changes, the full-length atomic model based on the Ca^2+^-bound structure (PDB ID: 4USL) was used as the initial model after removal of all Ca^2+^ ions. To study Sorcin activation and Ca^2+^-binding induced conformational changes, the full-length atomic model based on the apo structure (PDB ID: 4UPG) was used as the initial model. Three Ca^2+^ ions per monomer were placed at the EF-hands 1-3 binding sites by locally aligning each EF-hand motif in the Ca^2+^-bound structure to the corresponding regions in the apo structure and transferring the Ca^2+^ coordinates accordingly.

### Preparation of the equilibrated systems using VMD and NAMD

Each atomic model of dimeric Sorcin was solvated in a cubic water box (150 Å × 150 Å × 150 Å) using TIP3P water molecules(*29*) and neutralized with KCl to a final concentration of 150 mM using VMD(*26*). The fully solvated and ionized systems contained approximately 325,000 atoms. Details of all nine simulations (SIM1-SIM9) are provided in Table 1.

**Table 1.** Summary of molecular dynamics simulations. The active conformation corresponds to the Ca^2+^-bound crystal structure (PDB ID: 4USL), whereas the inactive conformation corresponds to the Ca^2+^-free (apo) crystal structure (PDB ID: 4UPG). Systems without Ca^2+^ occupancy were generated either from the apo structure or by removing Ca^2+^ ions from the Ca^2+^-bound structure. Conversely, Ca^2+^-occupied systems were prepared either by retaining Ca^2+^ ions in the Ca^2+^-bound structure or by introducing Ca^2+^ ions into the apo structure. “Minimal” Ca^2+^ restraints indicates that each Ca^2+^ ion was restrained to a single negatively charged coordinating residue within each EF-hand (Glu53, Glu94, and Glu124 for EF-hands 1, 2, and 3, respectively), whereas “extensive” Ca^2+^ restraints indicates that Ca^2+^ ions were restrained to all negatively charged coordinating residues within each EF-hand. All simulations were performed for 10 μs on Anton 3, except for SIM6, in which the first 0.4 μs were carried out using NAMD, followed by 9.6 μs on Anton 3. All MD trajectories were rendered as movies and shown in the Supporting Information.

| Simulation Name | Initial Conformation | Initial $\text{Ca}^{2+}$ occupancy | # of Atoms | $\text{Ca}^{2+}$ restrains | Spring Constant ($\text{kcal}\cdot\text{mol}^{-1}\cdot\text{\AA}^{-2}$) | Simulation length ($\mu\text{s}$) |
| --- | --- | --- | --- | --- | --- | --- |
| SIM1 | Active | No | 323,582 | - | - | 10 |
| SIM2 | Active | Yes | 324,637 | No | - | 10 |
| SIM3 | Active | Yes | 324,637 | minimal | 0.05 | 10 |
| SIM4 | Active | Yes | 324,637 | extensive | 0.05 | 10 |
| SIM5 | Active | Yes | 324,637 | extensive | 1 | 10 |
| SIM6 | Inactive | No | 320,504 | - | - | 10 |
| SIM7 | Inactive | Yes | 325,351 | No | - | 10 |
| SIM8 | Inactive | Yes | 325,351 | minimal | 0.05 | 10 |
| SIM9 | Inactive | Yes | 325,351 | extensive | 0.05 | 10 |

A fully equilibrated system is required to initiate production simulations on Anton 3. This was achieved through energy minimization and equilibration using NAMD3(*30*) with the CHARMM36m force field(*28*). Each system was subjected to 5,000 steps of conjugate-gradient energy minimization, followed by 10 ns of MD simulations. During the first 1 ns, all heavy atoms of the protein and Ca^2+^ ions were harmonically restrained. This was followed by an additional 1 ns of simulation with only the protein backbone harmonically restrained. The final 8 ns consisted of standard equilibrium MD simulations without any restrains in the NPT ensemble at 1 atm.

The temperature was maintained at 310 K using a Langevin thermostat (damping constant of 5.0 ps^-1^), and the pressure was controlled using a Nosé-Hoover Langevin piston barostat(*31*). Long-range electrostatics were treated using the particle-mesh Ewald (PME) method(*32*) under periodic boundary conditions. The van der Waals cutoff was set to 12 Å, with a smoothing function applied from 10 to 12 Å. One of the nine simulations (apo Sorcin without Ca^2+^, SIM6 in Table 1) was extended for an additional 400 ns in the NVT ensemble at 310 K using a Langevin thermostat with a damping constant of 1.0 ps^-1^. This 400 ns trajectory was treated as part of the production run and continued on Anton 3 as described below.

### Long-time scale MD simulation on Anton 3

The production runs on Anton 3 were initiated from the last frame of the equilibration simulations from NAMD. The CHARMM36m force field and TIP3P water model were used(*28*). The simulations were performed in the NVT ensemble at 310 K using the Antithetic thermostat and the Multigrator framework(*33*). The simulation time step was 2.5 fs, using a modified r-RESPA integrator(*34*). The short-range forces were evaluated every time step and long-range electrostatics every three time steps. The electrostatic interactions were computed using the u-series/SinhGS method(*35*). The van der Waals cutoff interactions were calculated using a cutoff of 9 Å. The bond lengths to hydrogen atoms were constrained using an implementation of M-SHAKE(*36*). Trajectory frames were saved every 1.2 ns for non-water atoms and every 12 ns for all atoms, respectively. Under these conditions, the achieved performance on Anton 3 was approximately 80 µs/day.

To prevent Ca^2+^ dissociation from the EF-hands, group distance restraints were applied in five of the nine simulations (Table 1) using a harmonic potential. Specifically, the distances between each Ca^2+^ ion and the carboxylate carbon atoms at the termini of the side chains of coordinating negatively charged residues (Asp and Glu) within the EF-hands were restrained. Spring constants of 0.05 kcal·mol^-1^·Å^-2^ (weak) or 1 kcal·mol^-1^·Å^-2^ (strong) were employed. The equilibrium distances were set based on values measured from the Ca^2+^-bound crystal structure of Sorcin (PDB ID: 4USL).

### Data Analysis

#### C_α_-Root Mean Square Deviation (RMSD)

Global conformational deviations were quantified using C_α_-RMSD relative to reference crystal structures (apo: 4UPG; Ca^2+^-bound: 4USL)(*20*). For each trajectory, frames were aligned to the reference structure based on C_α_ atoms to remove overall translation and rotation prior to RMSD calculation.

#### Angle between helices D and G

To quantify CTD rearrangements, the angle between helices D (residues 92-112) and G (residues 157-176) was calculated for each monomer, as used in the original report of the crystal structures(*20*). Helical axes were defined using the principal axis of the mass-weighted moment of inertia tensor of backbone atoms and were oriented consistently with respect to the initial structure.

#### Ca^2+^ ion coordination and dissociation

Ca^2+^ binding stability and dissociation from EF-hands was monitored by measuring distances between Ca^2+^ ions and key coordination residues in EF-hands. Specifically, the distances between each Ca^2+^ ion and the carboxylate carbon atoms at the termini of the side chains of coordinating negatively charged residues (Asp and Glu) within the EF-hands were monitored over time.

## Results and Discussions

Sorcin functions as a Ca^2+^ sensor by responding to changes in cytosolic Ca^2+^ concentration. Elevated Ca^2+^ levels promote Ca^2+^ binding to Sorcin, triggering conformational changes that expose hydrophobic surfaces(*20*). This Ca^2+^-bound conformation represents the active state, which can interact with target proteins such as calcium channels and pumps. Conversely, the Ca^2+^-free (apo) conformation corresponds to an inactive state that is less likely to engage in protein-protein interactions. Both the Ca^2+^-bound active state and the apo inactive state have been previously resolved by X-ray crystallography(*20*). The structural differences between these states are well defined. The goal of our long-timescale MD simulations is to elucidate the Ca^2+^-induced structural transitions of Sorcin between these active and inactive states, as well as to characterize the structural ensembles associated with each state.

### Structural transition of Sorcin from the active to inactive state

To investigate how Ca^2+^ release induces the transition of Sorcin from the active to the inactive state, we performed a 10 μs MD simulation starting from the Ca^2+^-bound active structure (PDB ID: 4USL) after removing all bound Ca^2+^ ions (SIM1 in Table 1). Structural convergence toward the apo inactive state was quantified by measuring the angle between helices D and G, a metric previously identified as the primary global conformational change associated with Ca^2+^ binding and release(*20*). In crystal structures, this angle shifts from 57° in the apo inactive state to 78° in the Ca^2+^-bound active state (Figure 1B and 1C). This change reflects a rotation of the EF1-EF2 subdomain away from the EF3-EF4-EF5 subdomain upon Ca^2+^ binding, thereby exposing additional hydrophobic surface(*20*).

Our simulation captures this transition from the active to the inactive state for chain A. The helix D-G angle decreased from ∼78° to as low as 54°, as shown in Figure 2A. Beyond resolving the continuum of transient intermediate states at atomic resolution, the simulation reveals an ultrafast structural response of Sorcin to Ca^2+^ release on the microsecond timescale. Such rapid responsiveness is likely critical for Sorcin’s function as a Ca^2+^ sensor, enabling it to quickly adapt to decreasing cytosolic Ca^2+^ levels, adopt the inactive conformation, and dissociate from binding partners.

**Figure 2.**
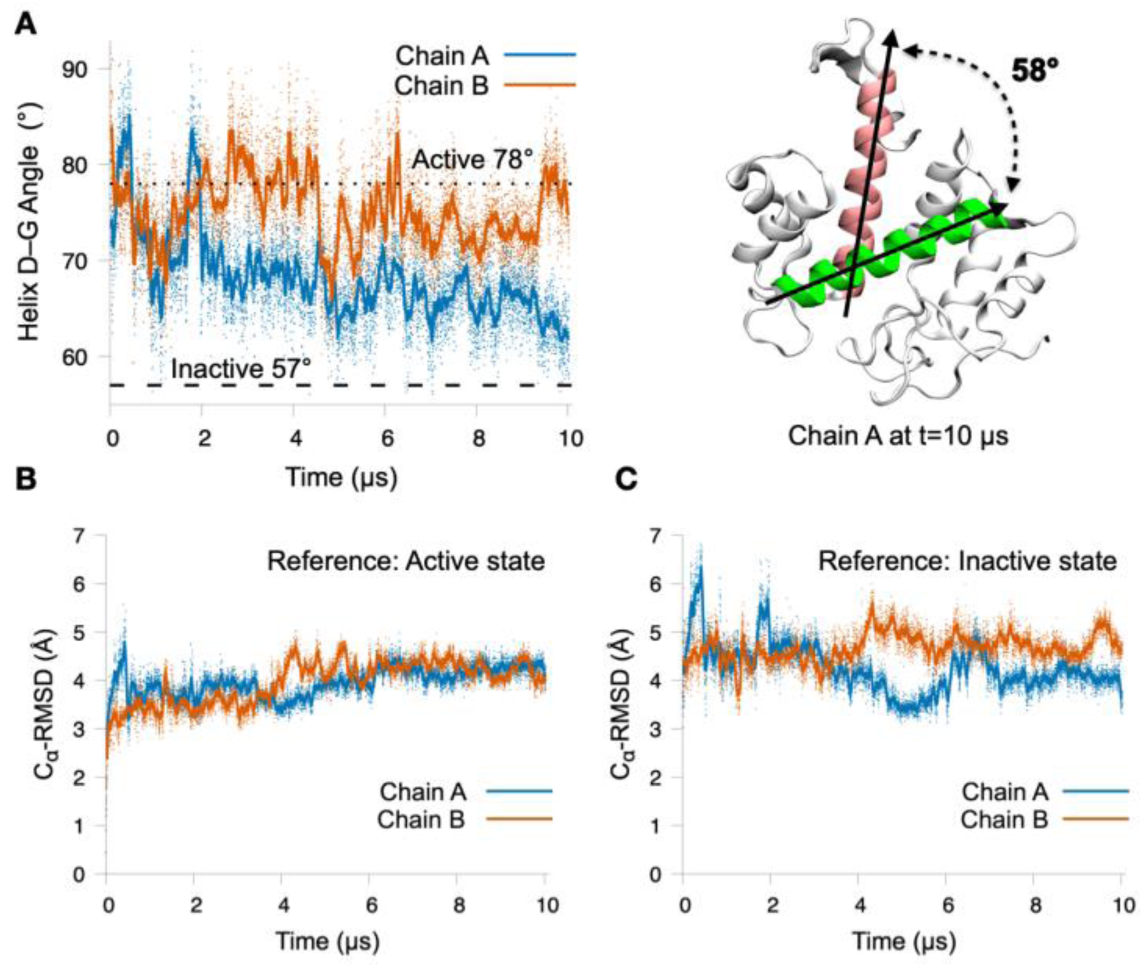
Structural transition following removal of Ca^2+^ from the Ca^2+^-bound active Sorcin structure (PDB ID: 4USL; SIM1 in Table 1). (A) Time evolution of the interhelical angle between helices D and G for Chain A (blue) and Chain B (orange). The angles corresponding to the Ca^2+^-bound active state (78°, PDB: 4USL) and the apo inactive state (57°, PDB ID: 4UPG) are indicated by dashed lines. The final conformation of Chain A closely resembles the apo inactive state and is shown on the right. (B) C_α_-RMSD of Chain A (blue) and Chain B (orange) relative to the Ca^2+^-bound active structure (PDB ID: 4USL) throughout the simulation. (C) C_α_-RMSD of Chain A (blue) and Chain B (orange) relative to the apo inactive structure (PDB ID: 4UPG) throughout the simulation.

Another indicator of structural convergence is the C_α_ root-mean-square deviation (C_α_-RMSD). Due to the intrinsic flexibility of the N-terminal domain (NTD), only residues resolved in the apo crystal structure (residues 30-198) were included in the RMSD analysis. This selection is further justified because all EF-hand motifs reside within the C-terminal domain (CTD, residues 34-198), where the Ca^2+^-induced global conformational changes occur. Following Ca^2+^ removal, Sorcin rapidly deviates from the initial active state, with the C_α_-RMSD reach higher than 4.5 Å within 10 μs (Figure 2B). Convergence toward the inactive state was assessed by calculating the C_α_-RMSD relative to the apo crystal structure (PDB ID: 4UPG). Although the simulation does not sample conformations identical to the apo structure (C_α_-RMSD = 0 Å), it reaches states with C_α_-RMSD values below 3.5 Å (Figure 2C). This remaining deviation likely reflects the intrinsic flexibility of the apo state, which will be further evaluated using simulation initiated from the apo crystal structure (SIM6 in Table 1). Supporting this interpretation, two apo crystal structures of human Sorcin obtained under different crystallization conditions (PDB ID: 4UPG and 1JUO) exhibit a C_α_-RMSD difference of ∼3.0 Å, underscoring the conformational heterogeneity of apo Sorcin(*19,20*).

### Ultrafast Ca^2+^ Dissociation from Sorcin

After directly observing the transition from the active to the inactive state in our simulations, we sought to confirm that this transition is driven by Ca^2+^ dissociation rather than the intrinsic flexibility of Sorcin. To this end, we performed a 10 μs MD simulation starting from the Ca^2+^-bound active structure (PDB ID: 4USL) while retaining all bound Ca^2+^ ions (SIM2 in Table 1). If no transition to the inactive state were observed in this control simulation, it would support the conclusion that the transition seen in SIM1 is specifically induced by Ca^2+^ dissociation.

Unexpectedly, we observed ultrafast Ca^2+^ dissociation from all EF-hand sites in this simulation (SIM2). As shown in Figure 3, the first dissociation events occurred on microsecond timescales. Specifically, the residence times before the first dissociation events were 4.5 and 0.8 μs for EF1, 0.5 and 1.0 μs for EF2, and 1.6 and 1.2 μs for EF3. The two residence times correspond to the two protomers in the Sorcin dimer. We also observed rapid rebinding of dissociated Ca^2+^ ions, either to their original EF-hand sites or to alternative EF hands. Because our analysis shown in Figure 3 monitors the distance between each Ca^2+^ ion and its initial binding site, stable rebinding to different EF hands appears as a relatively constant distance greater than 5 Å over timescales of hundreds of nanoseconds along the trajectories (Figure 3). The exact EF-hand site to which each Ca^2+^ ion rebinds was determined by monitoring the distances between the each Ca^2+^ ion and all EF-hand sites throughout the simulation (see Supporting Information, Figure S1-3).

**Figure 3.**
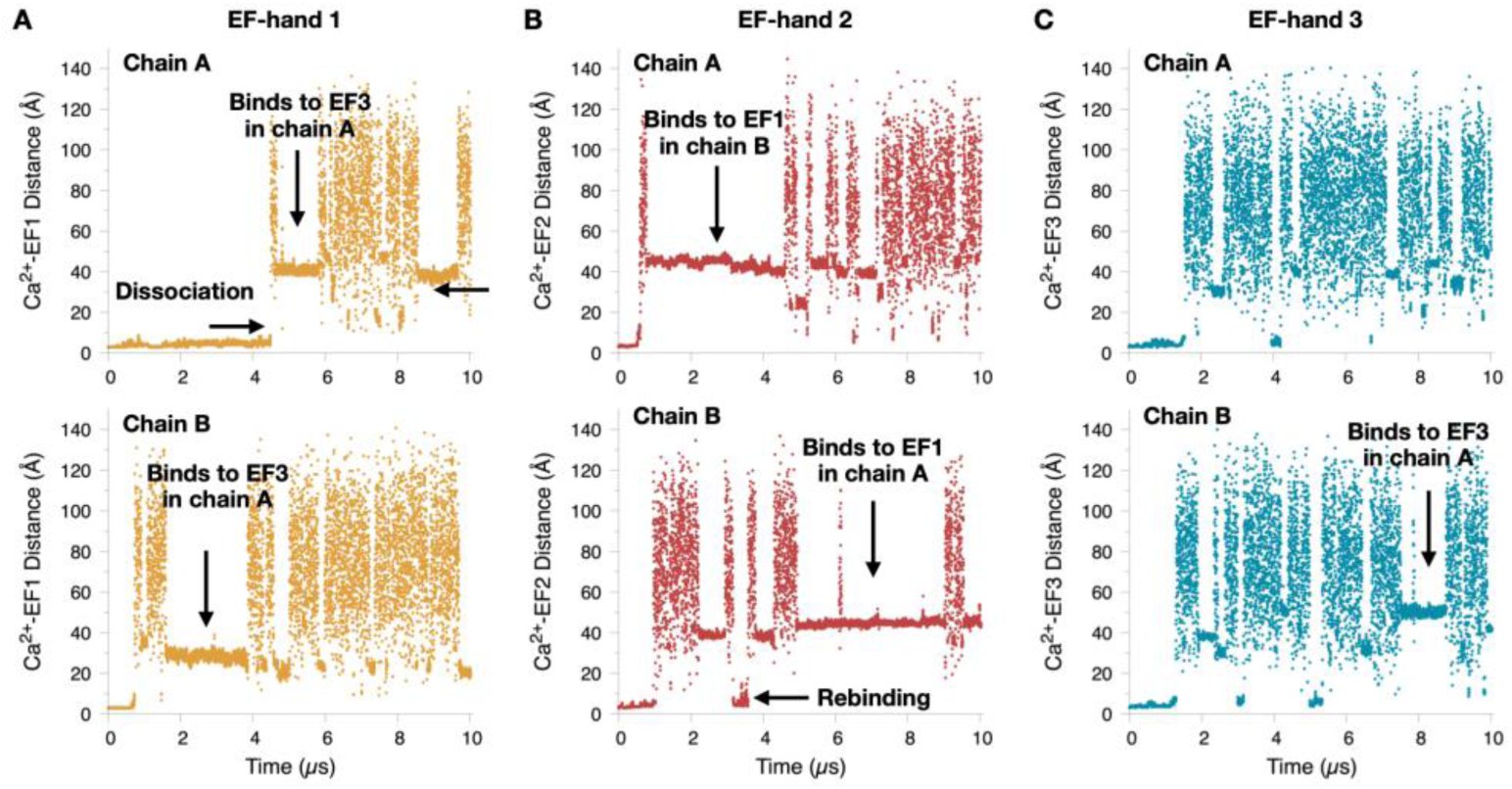
Spontaneous Ca^2+^ dissociation from and rebinding to EF-hand sites. (A) Ca^2+^ dissociation events from EF-hand 1 in Chain A (upper) and Chain B (lower). The dissociated Ca^2+^ ion subsequently rebinds to other EF-hand sites later in the simulation. (B) Ca^2+^ ions rapidly dissociate from EF-hand 2 in Chain A (upper) and Chain B (lower). In Chain B, reversible dissociation-rebinding-dissociation of the same Ca^2+^ ion was observed at EF-hand 2. (C) Ca^2+^ dissociation events from EF-hand 3 in Chain A (upper) and Chain B (lower). Selected rebinding events, including rebinding to the original EF-hand site or binding to a different EF-hand site, are indicated by arrows. The EF-hand site to which each Ca^2+^ ion rebinds was determined by monitoring the distances between the Ca^2+^ ion and all EF-hand sites throughout the simulation (see Supporting Information, Figure S1).

Although SIM2 did not function as the intended control simulation, it reveals remarkably fast binding and unbinding kinetics of Ca^2+^ ions at Sorcin EF-hand sites. In principle, the association and dissociation rate constants (k_on_ and k_off_) can be computed directly from simulation trajectories if enough reversible binding and dissociation events were observed. However, due to limited sampling, we provide only qualitative estimates. The residence time of Ca^2+^ is on the order of microseconds, corresponding to k_off_ ∼10^6^ s^-1^, indicative of extremely rapid dissociation. For comparison, the well-studied Ca^2+^ binding protein Calmodulin exhibits k_off_ values of ∼300-500 s^-1^ for its N-lobe EF hands(*37–39*). Note that binding of active Sorcin to its target proteins may dramatically change the residence time of Ca^2+^ and the k_off_.

Estimating k_on_ is more challenging due to its dependence on the free Ca^2+^ concentration, which is not constant in our simulations. Initially, no free Ca^2+^ ions are present. However, each dissociation event releases one Ca^2+^ ion into the solvent, resulting in an approximate increase of ∼0.5 mM in free Ca^2+^ concentration given the relatively small simulation box. Consequently, the free Ca^2+^ concentration fluctuates dynamically as ions dissociate and rebind to the protein surface or EF-hand sites. Assuming an effective free Ca^2+^ concentration in the millimolar range and a diffusion-limited rebinding timescale on the order of microseconds, we estimate k_on_ ∼10^9^ M^-1^ s^-1^. This rough estimate yields a dissociation constant (K_d_) on the order of millimolar, compared to an apparent experimental value of ∼0.1 mM obtained from our stopped-flow aggregation assays(*40*).

The discrepancy likely reflects both limited sampling and known limitations of classical force fields in accurately modeling divalent cations such as Ca^2+^(*41,42*). More extensive simulations and improved parameterization such as in the polarizable force field will be required for quantitative determination of these kinetic parameters at individual EF-hand sites(*43,44*). Nonetheless, the observed ultrafast binding and unbinding kinetics are consistent with Sorcin’s role as a Ca^2+^ sensor. Both the ion exchange dynamics and the conformational response to Ca^2+^ release occur on rapid timescales, suggesting that Sorcin can respond to fluctuations in intracellular Ca^2+^ concentration without a rate-limiting step.

### Weakly Restrained Ca^2+^ Coordination Reveals Intrinsic Instability of EF-Hand Binding

To prevent Ca^2+^ dissociation from the EF-hand sites, we performed a 10 μs simulation with distance restraints (SIM3 in Table 1). A single distance restraint was applied in each EF hand between the Ca^2+^ ion and the bidentate glutamate residues (Glu53, Glu94, and Glu124 in EF1, EF2, and EF3, respectively). These glutamate residues were selected because they provide two oxygen atoms for coordination and thus represent the most stable Ca^2+^-ligand interactions within each EF hand(*20*). To minimize perturbations introduced by the restraints, only one coordination interaction per EF hand was restrained. In addition, a weak harmonic restraint with a spring constant of 0.05 kcal·mol^-1^·Å^-2^ was used to further reduce artificial bias.

The distances between Ca^2+^ ions and negatively charged residues in the EF loops are shown in Figure 4. As expected, Ca^2+^ ions remained coordinated to the bidentate glutamates throughout the simulations. However, transient excursions were still observed, with Ca^2+^-Glu distances occasionally exceeding 5 Å. This behavior likely reflects the weak restraint strength and highlights the intrinsic instability of Ca^2+^ binding, even at these bidentate sites. Other coordinating residues, primarily aspartates, exhibited even greater instability. For example, coordination between Ca^2+^ and Asp46 in EF1 repeatedly broke and reformed during the simulation, with distances occasionally exceeding 15 Å. Similarly, coordination with Asp85 in EF2 was lost near the end of the simulation in both protomers, with distances exceeding 20 Å. In contrast, EF3 displayed the most stable coordination, with Ca^2+^-Asp distances remaining below 10 Å for the majority of the trajectory.

**Figure 4.**
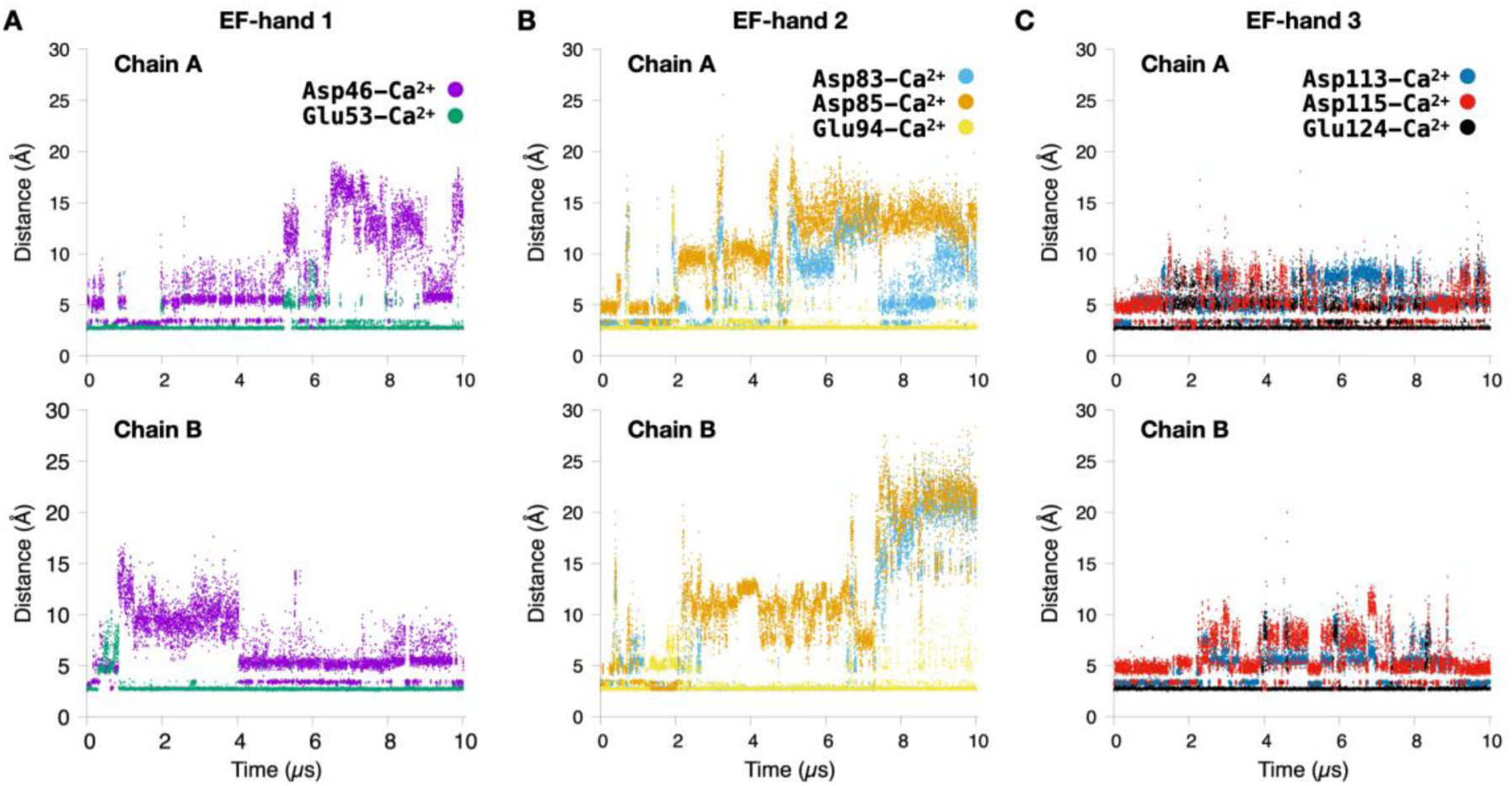
Distances between Ca^2+^ ions and negatively charged coordinating residues in EF-hands during simulations with weak distance restraints (force constant, k = 0.05 kcal·mol^-1^·Å^-2^) applied to the bidentate glutamate ligand. (A) Distances between Ca^2+^ and the coordinating residues Asp46 and Glu53 of EF-hand 1 in Chain A (upper) and Chain B (lower). The Ca^2+^-Glu53 distance was weakly restrained. (B) Distances between Ca^2+^ and the coordinating residues Asp83, Asp85, and Glu94 of EF-hand 2 in Chain A (upper) and Chain B (lower). The Ca^2+^-Glu94 distance was weakly restrained. (C) Distances between Ca^2+^ and the coordinating residues Asp113, Asp115, and Glu124 of EF-hand 3 in Chain A (upper) and Chain B (lower). The Ca^2+^-Glu124 distance was weakly restrained.

Importantly, all fluctuations in Ca^2+^-Asp distances occurred while Ca^2+^ remained restrained to the bidentate glutamate and localized within the EF-hand binding region. These observations further underscore the inherently weak binding affinity of Ca^2+^ to Sorcin EF-hand sites. Due to the partial loss of Ca^2+^ coordination over the course of the trajectory, this simulation likely represents intermediate states along the Ca^2+^ binding or release pathway and therefore cannot be used to reliably characterize the conformational ensemble of the fully Ca^2+^-bound active state.

### Conformational ensemble of Sorcin in the Ca^2+^-bound active state

We are particularly interested in simulations of the Ca^2+^-bound active state. Beyond serving as controls to support the conclusion that the transition observed in SIM1 is induced by Ca^2+^ release, these simulations provide critical insight into the conformational ensemble of Sorcin in its Ca^2+^-bound active form. To address the weak binding affinity of Ca^2+^ to EF-hand sites, we next applied more extensive harmonic distance restraints than in SIM3, restraining Ca^2+^ ions to all negatively charged coordinating residues within each EF hand. Two independent 10 μs simulations were performed using these restraint schemes. The two simulations differ in restraint strength, with spring constants of 0.05 and 1 kcal·mol^-1^·Å^-2^ applied in SIM4 and SIM5, respectively.

Using these refined restraint schemes, Ca^2+^ ions were retained within the EF-hand binding sites throughout the simulations. However, substantial fluctuations in the distances between Ca^2+^ ions and their coordinating residues were still observed in the weakly restrained SIM4, particularly for the aspartate ligands (Figure S4). Increasing the restraint strength to a force constant of 1 kcal·mol^-1^·Å^-2^ in SIM5 markedly reduced these distance fluctuations, resulting in more stable Ca^2+^ coordination (Figure S5). The resulting conformational ensembles were characterized using two-dimensional heat maps constructed from the C_α_-RMSD relative to the apo inactive state (PDB ID: 4UPG) and the angle between helices D and G as reaction coordinates. This analysis was performed for all five simulations (SIM1-5) initiated from the Ca^2+^-bound active state and is summarized in Figure 5 and Figure S6. In these plots, the inactive state is defined by a helix D-G angle of ∼57° and a C_α_-RMSD below 3.5 Å relative to the apo crystal structure.

**Figure 5.**
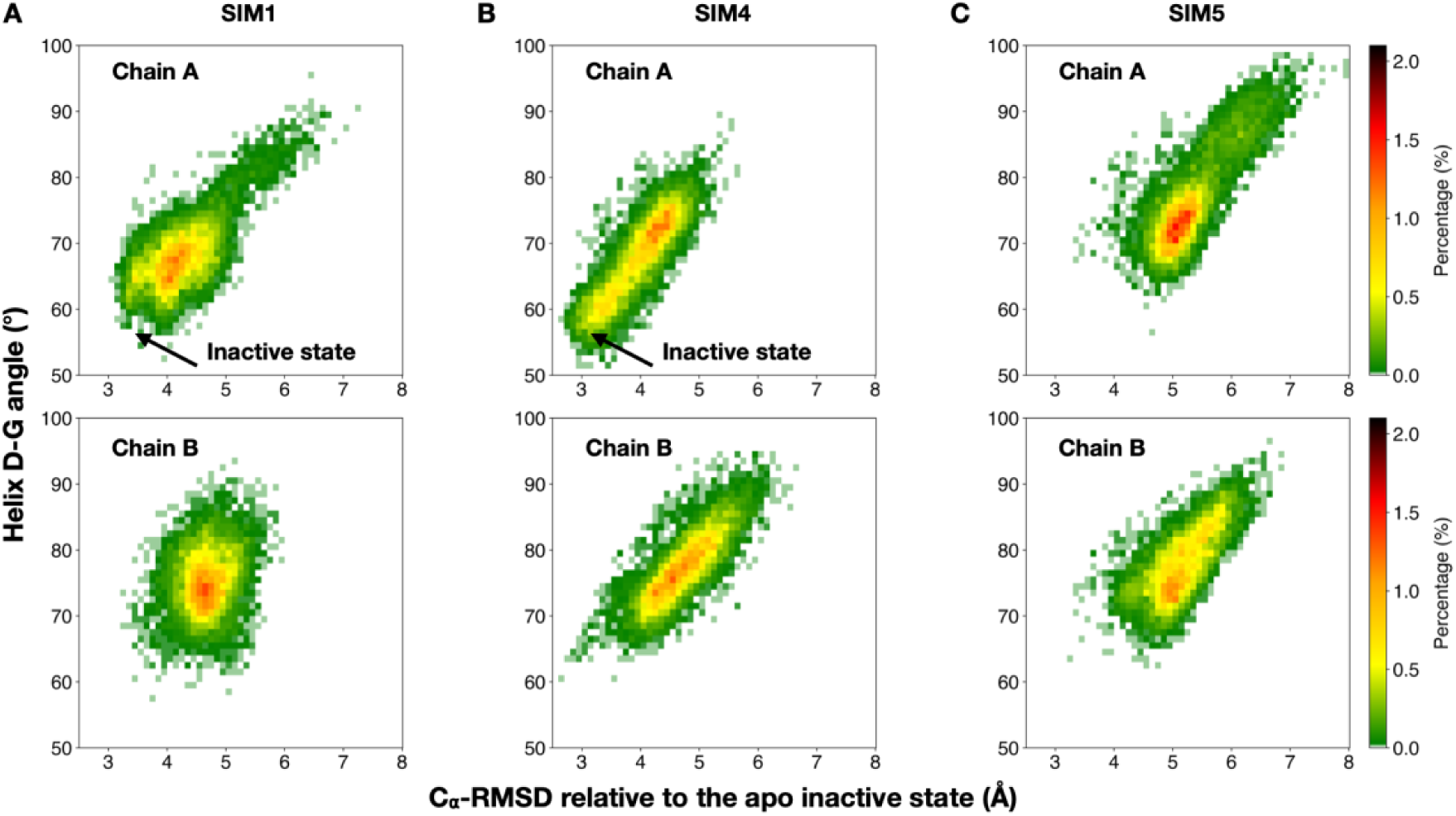
Heat maps of the conformational ensembles sampled in simulations initiated from the active state. (A) Conformational ensemble from SIM1, in which Ca^2+^ was removed from all EF-hand sites. Chain A transitioned to and sampled the inactive state. (B) Conformational ensemble from SIM4, in which weak distance restraints (force constant, k = 0.05 kcal·mol^-1^·Å^-2^) were applied between Ca^2+^ and the negatively charged coordinating residues of EF-hands 1-3. Chain A sampled the inactive state. (C) Conformational ensemble from SIM5, in which strong distance restraints (k = 1 kcal·mol^-1^·Å^-2^) were applied between Ca^2+^ and the negatively charged coordinating residues of EF-hands 1-3. Both Sorcin chains remained in the active state throughout the 10-μs simulation.

As discussed above, both protomers in SIM1, in which Ca^2+^ ions were removed at the beginning of the simulation, transitioned to the inactive state (Figure 5A). Similarly, both protomers in SIM2, where Ca^2+^ ions were present but not restrained, also reached the inactive state (Figure S6A). This outcome is expected, as Ca^2+^ ions dissociated from the EF-hands on the microsecond timescale, rendering SIM2 effectively similar to SIM1, despite occasional rebinding events. As shown previously, the weak single distance restraint applied in SIM3 was insufficient to maintain Ca^2+^ coordination within the EF-hands, leading chain A to sample the inactive state (Figure S6B). In SIM4, the application of more extensive but still relatively weak restraints stabilized protomer B in the active state, while protomer A continued to sample the inactive conformation (Figure 5B). In contrast, the stronger and extensive restraints applied in SIM5 successfully maintained both protomers in the active state throughout the 10 us simulation (Figure 5C). Together, these results support the conclusion that the active conformation of Sorcin is stabilized by Ca^2+^ binding to the EF-hands, and that the transition from the active to inactive state is driven by loss of coordination or Ca^2+^ release.

### Spontaneous Sampling of the Active State from Simulation of Apo State

After establishing that the conformational ensemble of the Ca^2+^-bound active state does not sample the inactive state, we next asked whether the Ca^2+^-free (apo) inactive ensemble can access the active conformation. To address this question, we performed a 10 μs simulation starting from the apo inactive structure of Sorcin (PDB ID: 4UPG) in the absence of Ca^2+^ (SIM6 in Table 1). In this simulation, the helix D-G angles of the two protomers evolved in opposite directions from the initial inactive value of ∼57° (Figure 6A). In protomer A, the angle decreased further to 30-40°, likely reflecting intrinsic flexibility of the apo state not captured in the crystal structure. In contrast, the same angle in protomer B increased to ∼78°, consistent with the active conformation.

**Figure 6.**
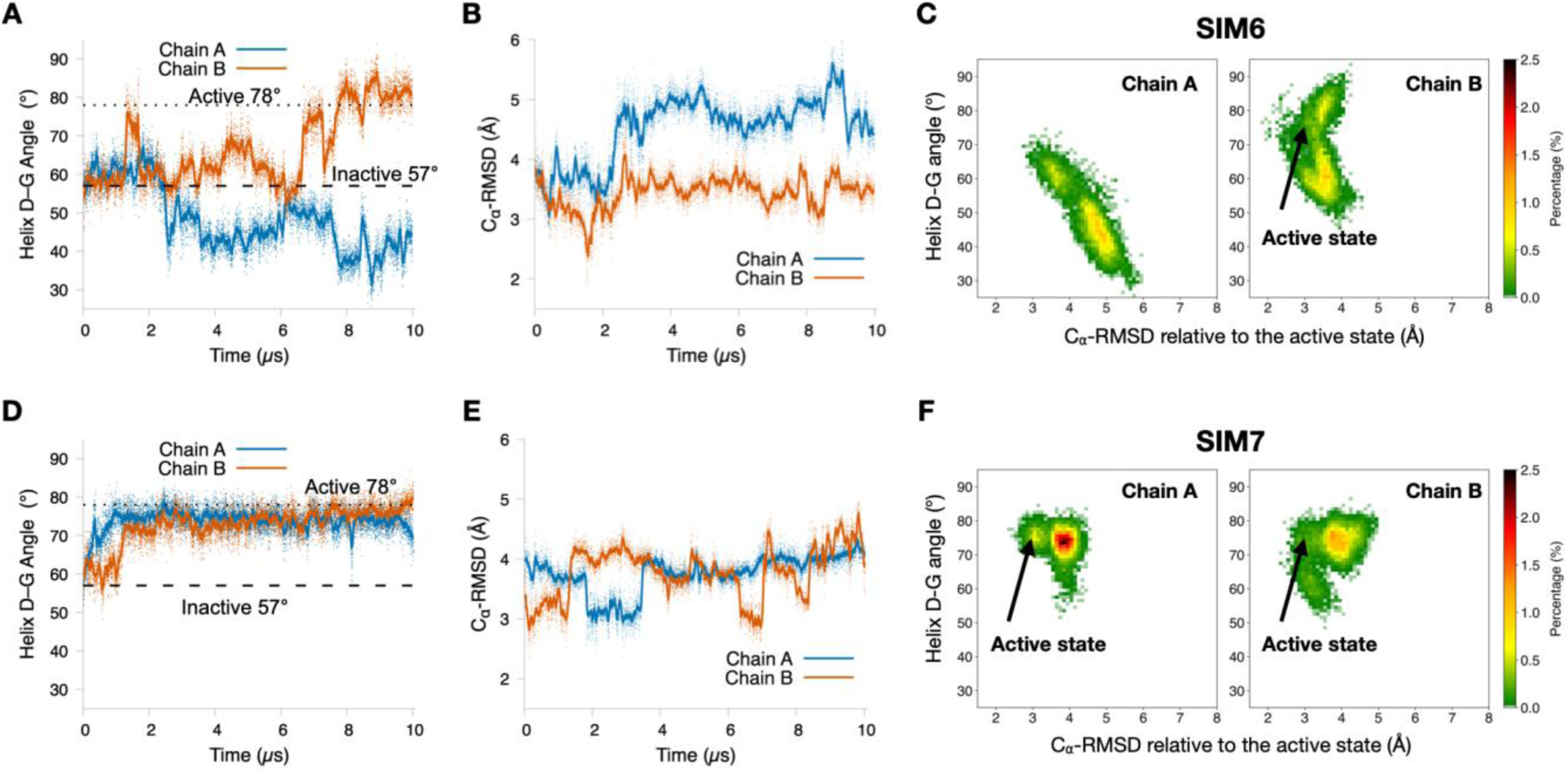
Structural transitions from the inactive to active states in the absence and presence of Ca^2+^. (A) Time evolution of the interhelical angle between helices D and G for Chain A (blue) and Chain B (orange) in SIM6, which was initiated from the apo inactive state in the absence of Ca^2+^. The interhelical angles of the Ca^2+^-bound active state (78°, PDB ID: 4USL) and the apo inactive state (57°, PDB ID: 4UPG) are indicated by dashed lines. (B) C_α_-RMSD of Chain A (blue) and Chain B (orange) relative to the Ca^2+^-bound active structure (PDB ID: 4USL) throughout SIM6. (C) Heat map of the conformational ensemble sampled during SIM6. The active-state conformations sampled by Chain B are indicated by an arrow. (D) Time evolution of the interhelical angle between helices D and G for Chain A (blue) and Chain B (orange) in SIM7, which was initiated from the apo inactive state with Ca^2+^ initially bound to EF-hands 1-3. During the simulation, all Ca^2+^ ions rapidly dissociated from their original binding sites and rebound to different EF-hands. (E) C_α_-RMSD of Chain A (blue) and Chain B (orange) relative to the Ca^2+^-bound active structure (PDB ID: 4USL) throughout SIM7. (F) Heat map of the conformational ensemble sampled during SIM7.

Convergence to the active state in protomer B was further confirmed by monitoring the C_α_-RMSD relative to the Ca^2+^-bound crystal structure (PDB ID: 4USL), which consistently dropped below 3 Å (Figure 6B). This active-like conformation was readily identifiable in the two-dimensional heat maps defined by the helix D-G angle and C_α_-RMSD relative to Ca^2+^-bound crystal structure (Figure 6C). These results indicate that the inactive-to-active transition can occur even in the absence of Ca^2+^, suggesting a relatively low kinetic barrier between the two states. We therefore hypothesize that Ca^2+^ binding shifts the conformational equilibrium by biasing population toward the active state rather than inducing the transition.

To test this hypothesis, we introduced Ca^2+^ ions into the EF-hand binding sites of the inactive structure. Specifically, each EF hand in the Ca^2+^-bound crystal structure (PDB ID: 4USL) was locally aligned to its counterpart in the apo structure, and the corresponding Ca^2+^ coordinates were transferred, yielding a Ca^2+^-bound but inactive starting configuration. A 10 μs simulation of this system was then performed without restraints (SIM7 in Table 1). Consistent with the weak Ca^2+^ binding observed in SIM2 and SIM3, all Ca^2+^ ions rapidly dissociated from the EF-hands in this simulation. Although transient binding and unbinding events were observed, the system cannot be considered stably Ca^2+^-bound. Nevertheless, the simulation sampled conformations characteristic of the active state, with the helix D-G angle converging to ∼78° and the C_α_-RMSD relative to the Ca^2+^-bound crystal structure dropping below 3 Å (Figure 6D, E and F). Together, the results from SIM6 and SIM7 demonstrate that the active conformation can be readily sampled in the absence of stable Ca^2+^ binding, supporting a model in which Ca^2+^ primarily stabilizes and shifts the population toward pre-existing active-like states.

To stabilize Ca^2+^ binding in the inactive state of Sorcin, we applied distance restraints between Ca^2+^ ions and negatively charged coordinating residues within each EF-hand. To minimize perturbation and bias, we first restrained only the distance between Ca^2+^ and the bidentate glutamate residues, as in SIM3, and performed a 10 μs simulation (SIM8 in Table 1). Similar to SIM3, the unrestrained coordination interactions were unstable (Figure S7). The trajectory exhibited behavior comparable to the Ca^2+^-free simulation (SIM6), with the helix D-G angles of the two protomers evolving in opposite directions. Notably, protomer A sampled the active state, with the helix D-G angle reached 78° (Figure 7A) and the C_α_-RMSD relative to the Ca^2+^-bound crystal structure dropping below 3 Å within 1 μs (Figure 7B). The active conformation was clearly identified as a highly populated state in the angle-RMSD heat map (Figure 7C). However, because Ca^2+^ was not fully coordinated and dissociated from other coordinating aspartate residues, this simulation does not represent a fully bound state despite the spatial proximity of Ca^2+^ to the EF-hands.

**Figure 7.**
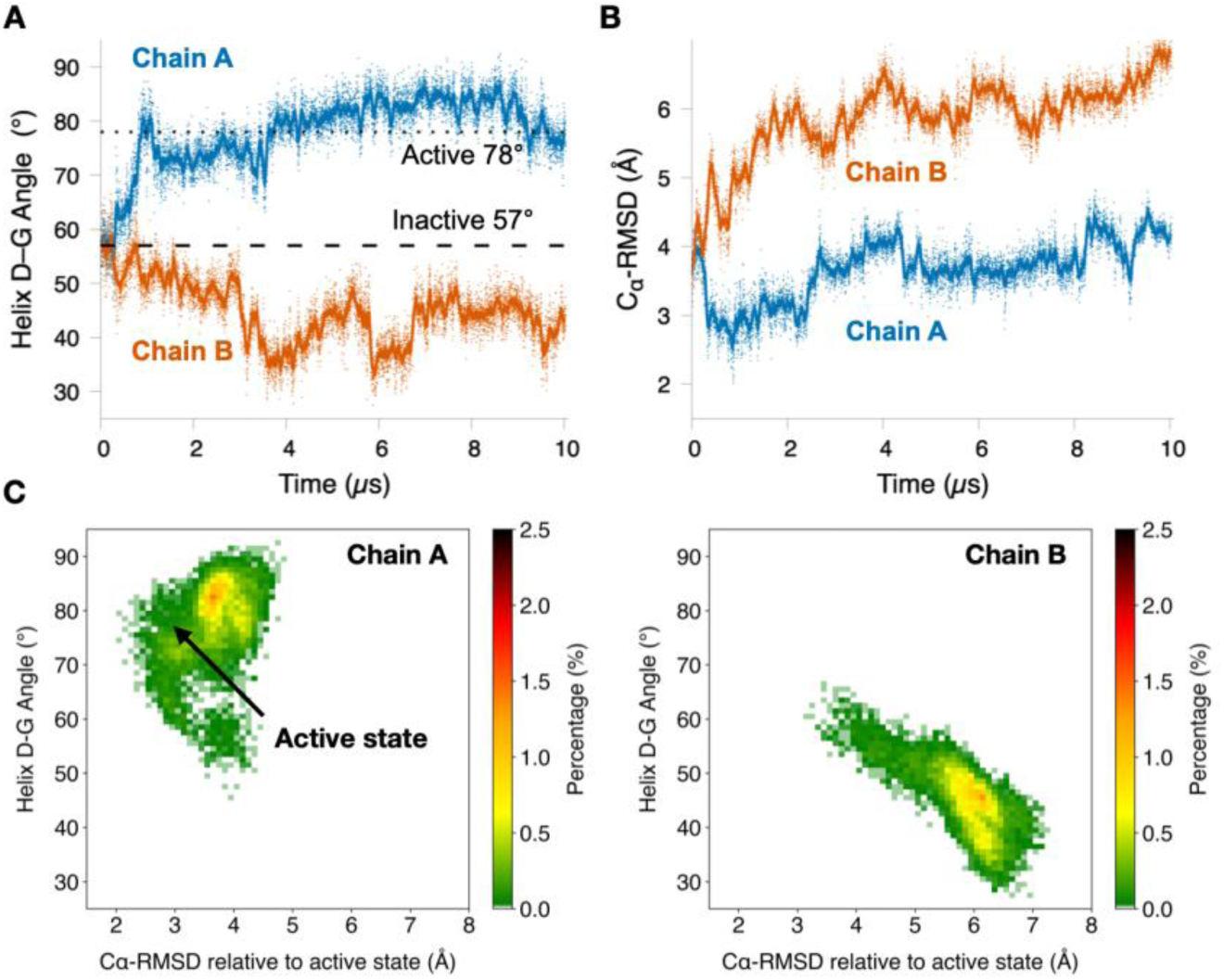
Structural transition toward the active state in SIM8 initiated from the apo inactive state with weak distance restraints (force constant, k = 0.05 kcal·mol^-1^·Å^-2^) applied between each Ca^2+^ ion and the bidentate glutamate ligand in its corresponding EF-hand. (A) Time evolution of the interhelical angle between helices D and G for Chain A (blue) and Chain B (orange). (B) C_α_-RMSD of Chain A (blue) and Chain B (orange) relative to the Ca^2+^-bound active structure (PDB ID: 4USL) throughout the simulation. (C) Heat map of the conformational ensemble sampled during SIM8. The active-state conformations sampled by Chain B are indicated by an arrow.

To model a fully coordinated Ca^2+^-bound inactive state, we next applied more extensive restraints, including distances between Ca^2+^ and all negatively charged coordinating residues in each EF-hand, as in SIM4, with a spring constant of 0.05 kcal·mol^-1^·Å^-2^, and performed a 10 μs simulation (SIM9 in Table 1). Surprisingly, this simulation did not sample the active state at all, with the helix D-G angles remaining below 70° for both protomers throughout the trajectory (Figure S8). Although unexpected, this result can be rationalized by a pathway-dependent transition model. If the inactive-to-active transition proceeds through specific intermediate states in a stepwise manner, imposing these restraints in the inactive conformation may trap Sorcin in an off-pathway state, thereby preventing progression toward the active state. Further work is needed to test this hypothesis.

### Structural Asymmetry in the Sorcin homodimer

Although all available crystal structures of Sorcin depict a symmetric homodimer(*19–21,45,46*), we consistently observe pronounced structural asymmetry in our simulations, with the C_α_-RMSD between the two protomers ranging from 2 to 8 Å (Figure 8). While crystal structures represent low-energy and spatially averaged conformations, MD simulations capture the full conformational ensemble near the native states, where asymmetry is expected even on short timescales. In the long-timescale simulations presented here, where transitions between inactive and active states occur in both directions, the degree of asymmetry becomes even more pronounced. In fact, we frequently observed in several simulations that one protomer completed a reversible transition between the inactive and active states, whereas the other protomer remained in a single conformational state throughout the simulation. This asymmetry may be further amplified by differences in Ca^2+^ occupancy at each EF hand between protomers or by distinct conformations of the flexible N-terminal domains (NTDs) on each protomer (not included in the current analysis). Such multilevel structural asymmetry could introduce an additional layer of regulation in Sorcin function, particularly in modulating its interactions with target proteins.

**Figure 8.**
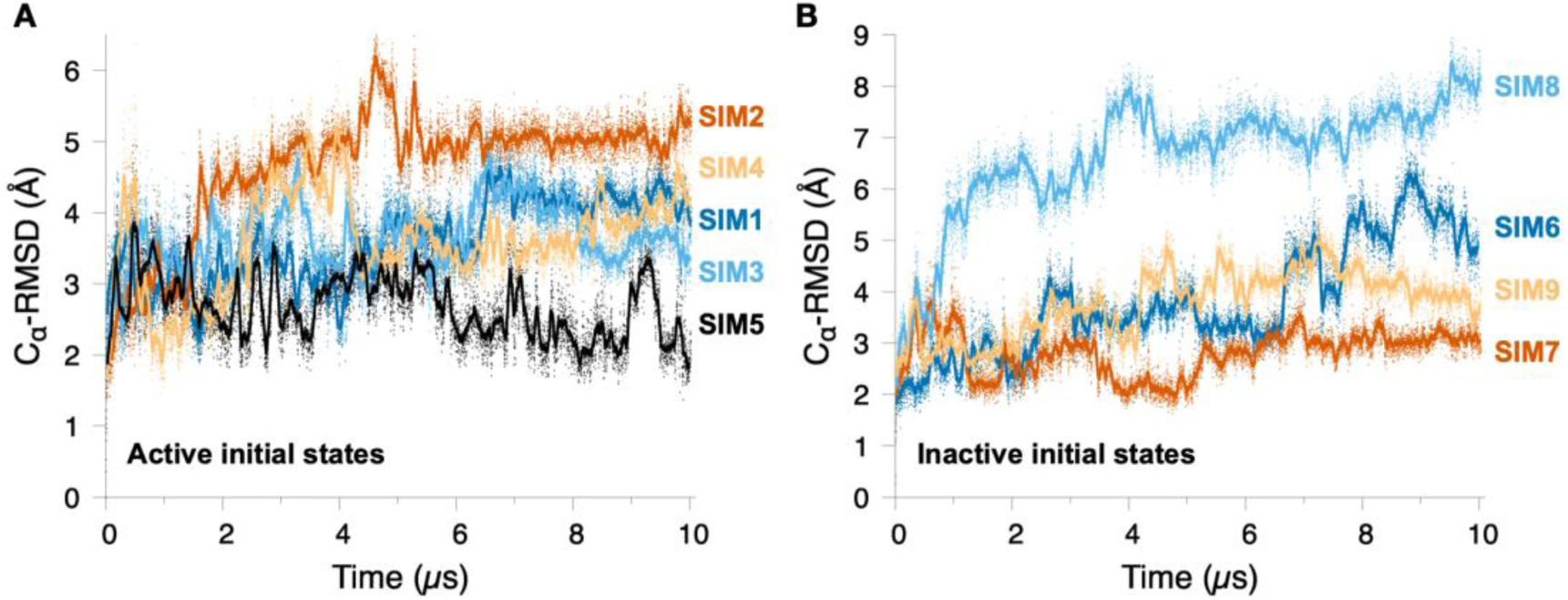
Structural asymmetry of the Sorcin dimer across all simulations. (A) Pairwise C_α_-RMSD between Chains A and B at each time point for simulations initiated from the active state. (B) Pairwise C_α_-RMSD between Chains A and B at each time point for simulations initiated from the inactive state. Simulation labels are defined in Table 1. Both structural alignment and C_α_-RMSD calculations were performed using residues 30-198, which are resolved in the crystal structures.

## Conclusions

In this work, we used long-timescale MD simulations to define the molecular mechanism by which Ca^2+^ regulates the conformational landscape of Sorcin. Our results provide a dynamic and mechanistic view that complements static structural data and leads to several key conclusions. First, we directly observed the transition from the Ca^2+^-bound active state to the apo inactive state following Ca^2+^ release, demonstrating that loss of Ca^2+^ coordination is sufficient to drive this conformational change. This transition occurs on the microsecond timescale, highlighting the rapid responsiveness of Sorcin as a Ca^2+^ sensor. Consistent with this behavior, we also observed ultrafast Ca^2+^ dissociation and rebinding events at EF-hand sites, indicating weak binding affinity and highly dynamic ion exchange. Second, simulations initiated from the apo inactive state reveal that Sorcin can spontaneously sample active-like conformations even in the absence of Ca^2+^. This finding indicates that the energy barrier between inactive and active states is relatively low and supports a conformational selection mechanism in which Ca^2+^ binding shifts the population toward pre-existing active states rather than inducing a new conformation. Finally, our simulations consistently reveal structural asymmetry between the two protomers of the Sorcin homodimer, in contrast to the symmetric crystal structures. This asymmetry is amplified during conformational transitions and may be further influenced by differential Ca^2+^ binding or N-terminal domain dynamics, suggesting a potential role in modulating interactions with target proteins. Overall, our results establish a unified model in which Sorcin functions through a combination of rapid Ca^2+^ exchange, conformational selection, and dynamic structural asymmetry. This work provides a comprehensive atomistic framework for understanding Sorcin activation and offers broader insights into the mechanisms of EF-hand Ca^2+^-binding proteins.

## Acknowledgments

We acknowledge the technical support provided by the Institute for Bioscience and Biotechnology Research (IBBR) Information Technology Department, particularly Gale Tempest and Scott Carlson. Y.L. and the Liu Laboratory are supported by start-up funding from the University of Maryland, College Park, and the State of Maryland. This project was also supported by an IBBR Seed Grant funded through the University of Maryland Strategic Partnership: MPowering the State. Anton 3 computer time was provided by the Pittsburgh Supercomputing Center (PSC) through Grant 1R24GM154042 from the National Institutes of Health. The Anton 3 machine at PSC is made available by D. E. Shaw Research. The authors also acknowledge the University of Maryland High Performance Computing (HPC) resources (https://hpcc.umd.edu), including the Zaratan HPC Cluster and the IBBR HPC Cluster, which were also used to conduct the research reported in this paper.

