## Supplemental Figure 1-8 for "Long-Timescale Molecular Dynamics Reveal a Coordination-Biased Conformational Selection Mechanism for Sorcin Activation"

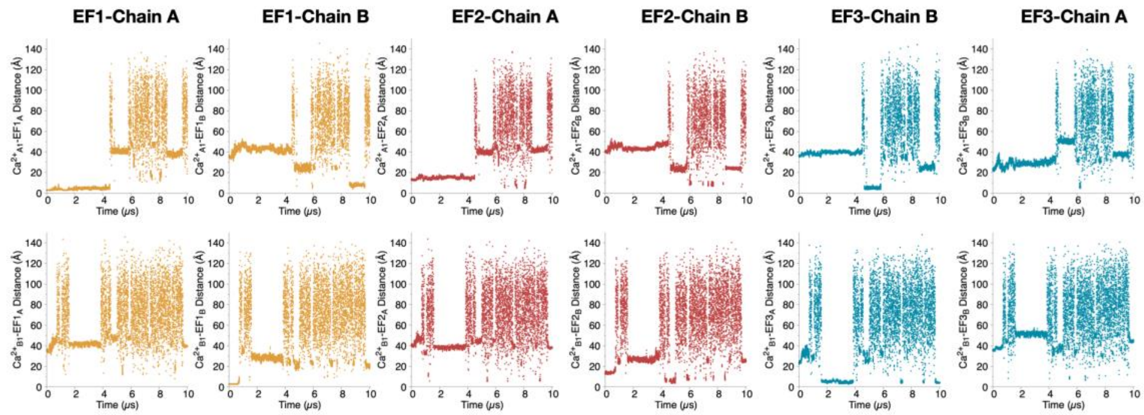


Figure S1. Distances between EF-hands 1-3 and each of the two $Ca^{2+}$ ions initially bound to EF-hand 1 throughout the simulation. These distance profiles were used to determine the EF-hand site occupied by each $Ca^{2+}$ ion at each time point or whether these $Ca^{2+}$ ions remained unbound in solution.

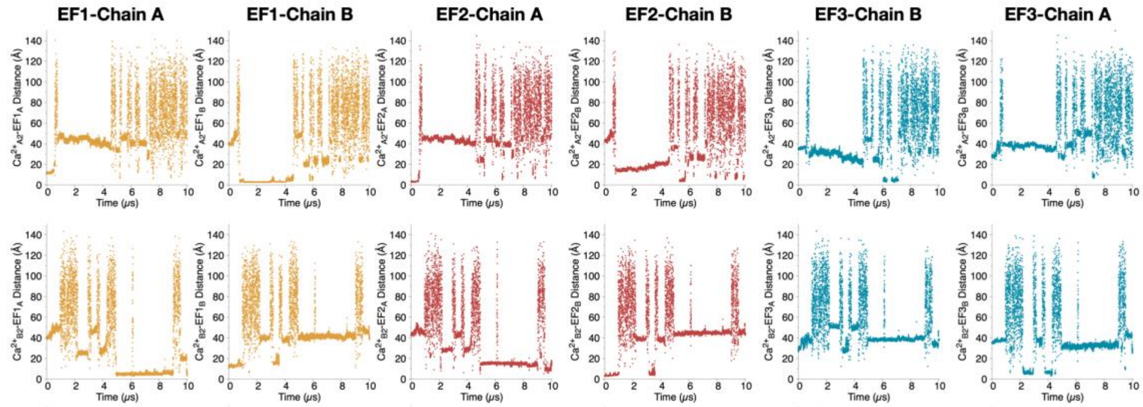


Figure S2. Distances between EF-hands 1-3 and each of the two $Ca^{2+}$ ions initially bound to EF-hand 2 throughout the simulation.

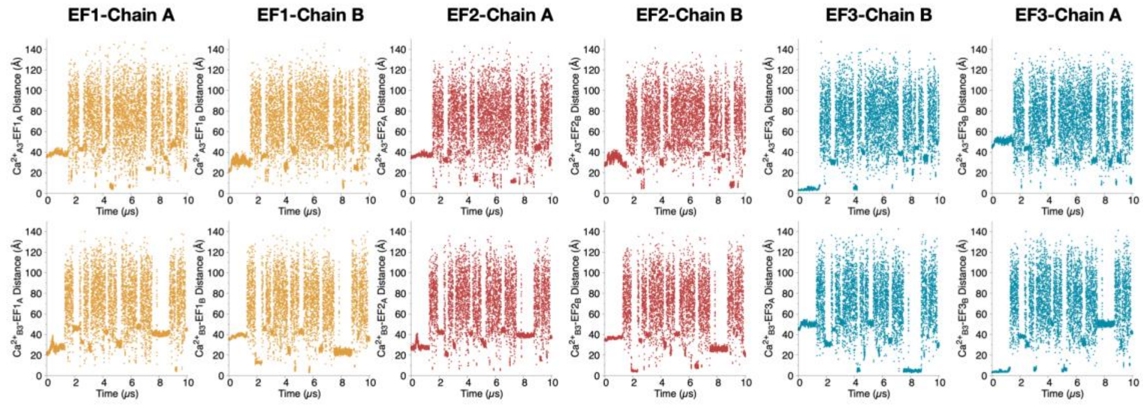


Figure S3. Distances between EF-hands 1-3 and each of the two $Ca^{2+}$ ions initially bound to EF-hand 3 throughout the simulation.

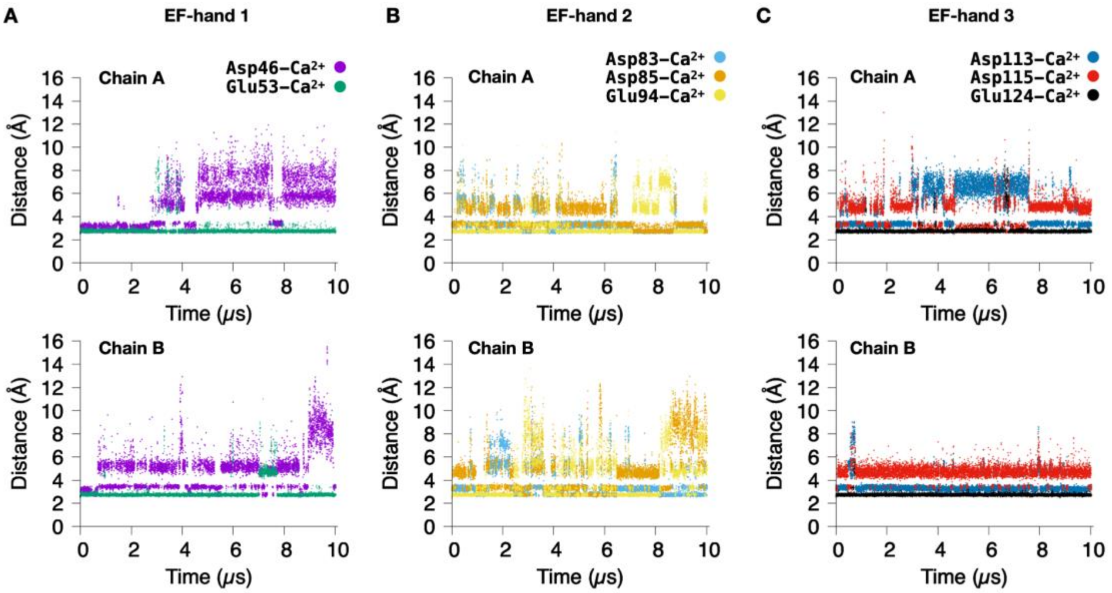


Figure S4. Distances between $Ca^{2+}$ ions and the negatively charged coordinating residues of the EF-hands during the SIM4 simulation, in which weak distance restraints (force constant, k = 0.05 kcal·mol$^{-1}$·Å$^{-2}$) were applied between $Ca^{2+}$ and all negatively charged coordinating residues in EF-hands 1-3. (A) Distances between $Ca^{2+}$ and the coordinating residues Asp46 and Glu53 of EF-hand 1 in Chain A (upper) and Chain B (lower). (B) Distances between $Ca^{2+}$ and the coordinating residues Asp83, Asp85, and Glu94 of EF-hand 2 in Chain A (upper) and Chain B (lower). (C) Distances between $Ca^{2+}$ and the coordinating residues Asp113, Asp115, and Glu124 of EF-hand 3 in Chain A (upper) and Chain B (lower).

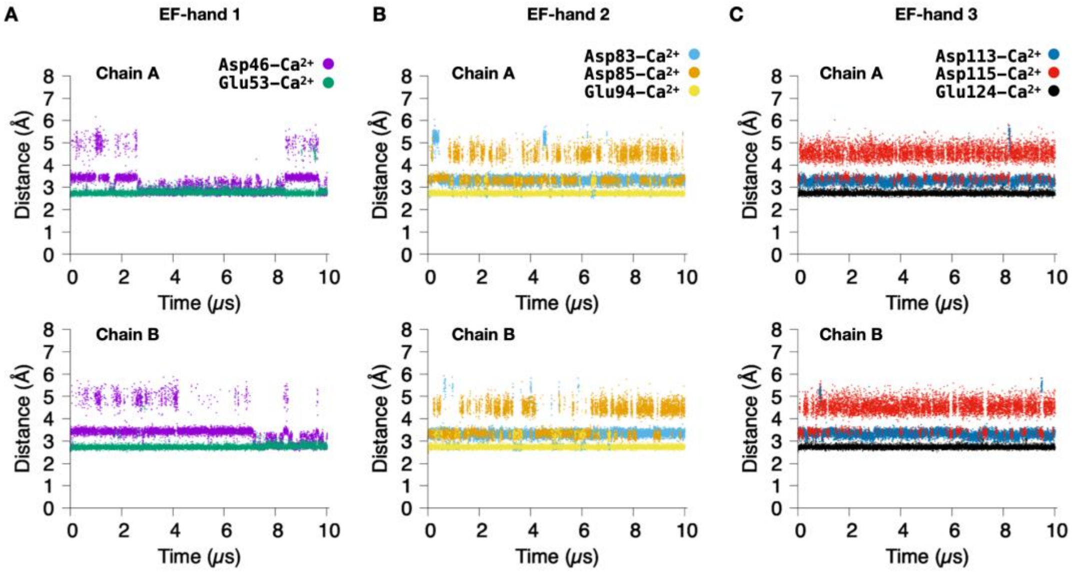


Figure S5. Distances between $Ca^{2+}$ ions and the negatively charged coordinating residues of the EF-hands during the SIM5 simulation, in which strong distance restraints (force constant, k = 1 kcal·mol$^{-1}$·Å$^{-2}$) were applied between $Ca^{2+}$ and all negatively charged coordinating residues in EF-hands 1-3. (A) Distances between $Ca^{2+}$ and the coordinating residues Asp46 and Glu53 of EF-hand 1 in Chain A (upper) and Chain B (lower). (B) Distances between $Ca^{2+}$ and the coordinating residues Asp83, Asp85, and Glu94 of EF-hand 2 in Chain A (upper) and Chain B (lower). (C) Distances between $Ca^{2+}$ and the coordinating residues Asp113, Asp115, and Glu124 of EF-hand 3 in Chain A (upper) and Chain B (lower).

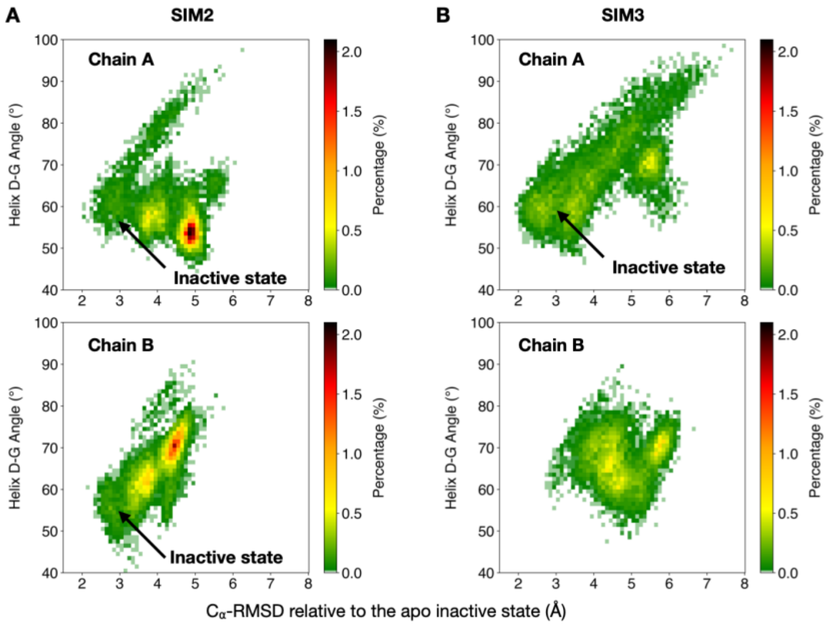


Figure S6. Heat maps of the conformational ensembles sampled in simulations initiated from the active state. (A) Conformational ensemble from SIM2, in which $Ca^{2+}$ ions were retained in all EF-hand sites without distance restraints. Both chains transitioned to and sampled the inactive state. (B) Conformational ensemble from SIM3, in which weak distance restraints (force constant, k = 0.05 kcal·mol$^{-1}$·Å$^{-2}$) were applied between $Ca^{2+}$ and the bidentate glutamate ligand of EF-hands 1-3. Chain A transitioned to and sampled the inactive state.

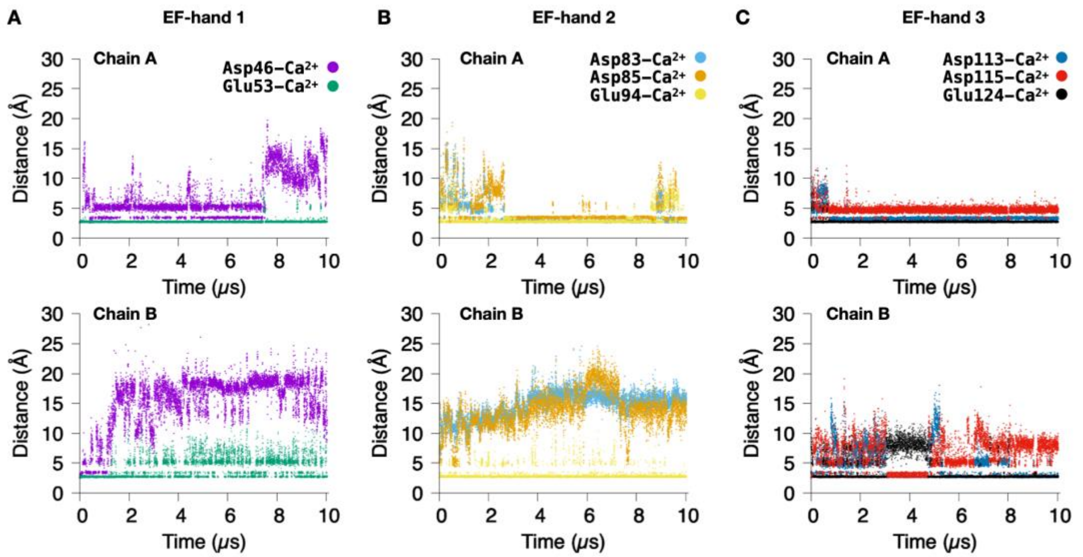


Figure S7. Distances between $Ca^{2+}$ ions and negatively charged coordinating residues in EF-hands in simulation SIM7 with weak distance restraints (force constant, k = 0.05 kcal·mol$^{-1}$·Å$^{-2}$) applied to the bidentate glutamate ligand. (A) Distances between $Ca^{2+}$ and the coordinating residues Asp46 and Glu53 of EF-hand 1 in Chain A (upper) and Chain B (lower). The $Ca^{2+}$-Glu53 distance was weakly restrained. (B) Distances between $Ca^{2+}$ and the coordinating residues Asp83, Asp85, and Glu94 of EF-hand 2 in Chain A (upper) and Chain B (lower). The $Ca^{2+}$-Glu94 distance was weakly restrained. (C) Distances between $Ca^{2+}$ and the coordinating residues Asp113, Asp115, and Glu124 of EF-hand 3 in Chain A (upper) and Chain B (lower). The $Ca^{2+}$-Glu124 distance was weakly restrained.

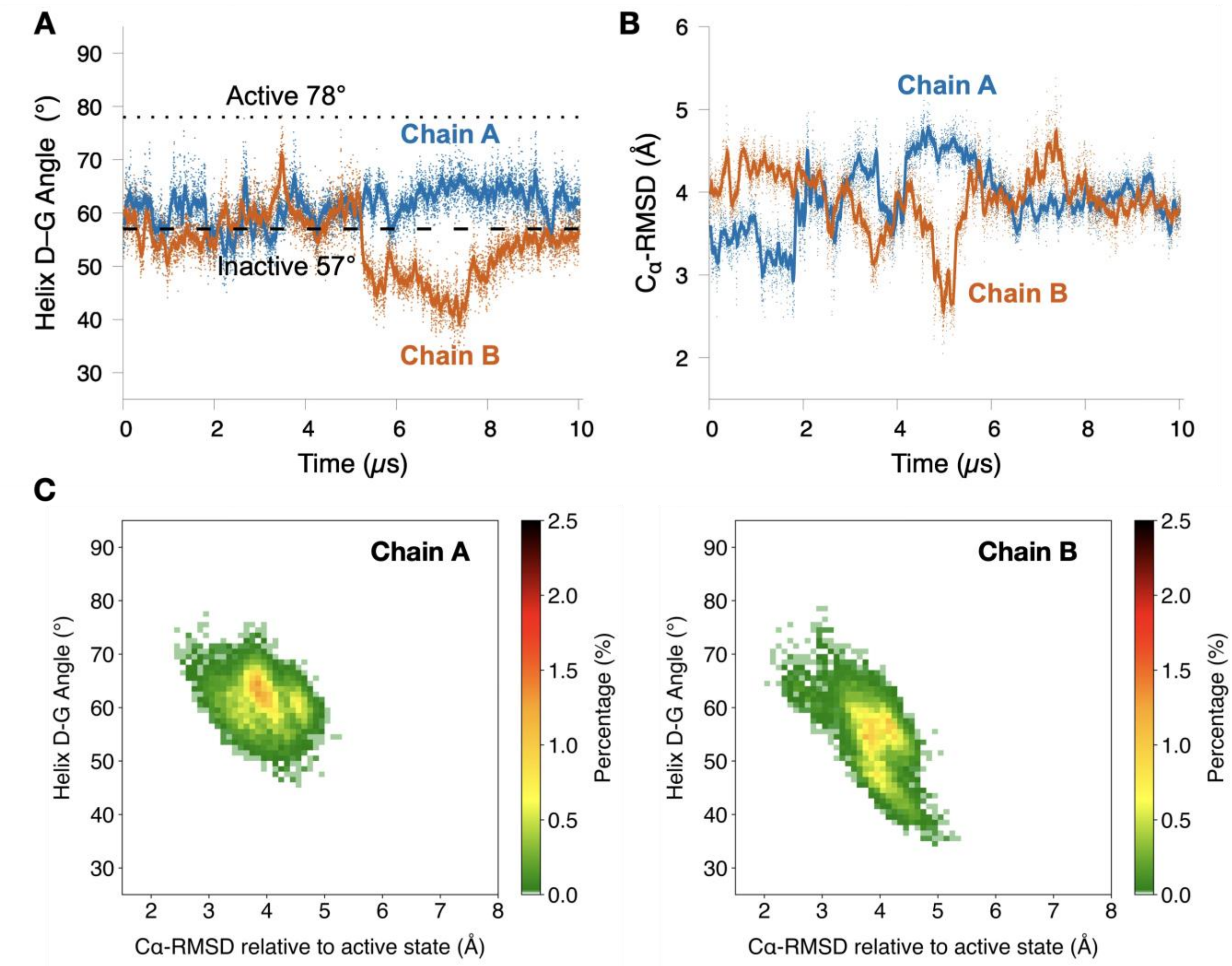


Figure S8. Conformational ensembles in simulation SIM9 which initiated from the apo inactive state with weak distance restraints (force constant, k = 0.05 kcal·mol$^{-1}$·Å$^{-2}$) applied between $Ca^{2+}$ and all negatively charged coordinating residues in EF hands 1-3. (A) Time evolution of the interhelical angle between helices D and G for Chain A (blue) and Chain B (orange). (B) $C_{\alpha}$-RMSD of Chain A (blue) and Chain B (orange) relative to the $Ca^{2+}$-bound active structure (PDB ID: 4USL). (C) Heat map of the conformational ensemble sampled.
